# An entropic measure of diverse specialisation highlights multifunctional neurons in annotated connectomes

**DOI:** 10.1101/2025.03.19.644231

**Authors:** Sung Soo Moon, Lidia Ripoll-Sánchez, Petra Vértes, William R. Schafer, Sebastian E. Ahnert

## Abstract

The creation and curation of synaptic-level neuronal networks, or connectomes, enables the study of the relationship between neuronal structure and function. Topological characteristics of neuronal networks have been studied extensively. Separately, there have been considerable efforts to classify the morphology, cell types, and lineages of neurons. Here, we introduce a network metric that combines topological analysis with node metadata. This entropic quantity measures the diversity of incoming or outgoing connections to a node in terms of the metadata distribution. We find that in *Caenorhabditis elegans*, the top-scoring neurons (`PVR, RMGL/R, DVA, CEP, ADE, URXR, RIGL, BAG, SMBDL`) have known functions that integrate and disseminate multi-modal information involved in sensorimotor functions. In the nerve cord of *Drosophila melanogaster*, we find that top-scoring neurons are embryonic neurons located in the abdominal neuropil, where sensorimotor coordination is required for complex innate behaviour such as mating.

## 1 Introduction

A brain is often modelled as a network [Bassett and Sporns, 2017, Bullmore and Sporns, 2009], but how are brain networks organised? Neuronal resolution synaptic mapping of the brain, or connectomes, are networks of nodes and weighted, directed edges between them. While the *C. elegans* connectome [White et al., 1986] was a landmark data set in neuroanatomy, we have yet to formulate a general network-level understanding of the organisational principles of nervous systems. More recently, the adult *Drosophila melanogaster* brain [Zheng et al., 2018, Scheffer et al., 2020, Dorkenwald et al., 2024] and nerve cord [Venkatasubramanian and Mann, 2019, Takemura et al., 2024, Cheong et al., 2024, Marin et al., 2024, Court et al., 2020, Stürner et al., 2024], data sets have been imaged, segmented and reconstructed, and annotated. With high-throughput experimental techniques granting more metadata about the nature of the neurons [Harris et al., 2015, Janssens et al., 2022] and the synaptic connections [Buhmann et al., 2021, Eckstein et al., 2024, Varshney et al., 2011, Cook et al., 2019], connectome data sets are complex annotated systems and not simple undirected networks. Annotations are key supplements to our understanding of brain structure and function, often related to dynamical properties of the system, and intrinsic to biological function [Bazinet et al., 2023a,b, Randi et al., 2023]. However, current methodologies often overlook them; extensions of network methodologies respecting annotation metadata in neuroscience are thus needed to bridge the gap between brain structure and function.

The application of network analysis to neuroscience has proved fruitful, establishing key interpretable features of brain networks, including small-worldness, modularity, hubs, rich-clubs and community structure [Bassett and Sporns, 2017, Bullmore and Sporns, 2009, Betzel et al., 2024, Towlson et al., 2013, Jarrell et al., 2012]. Many analyses focus on assigning an importance to a node in the context of global topology, like centrality measures. However, measures based on local topology may be more appropriate for studying biological networks with specific cross-modular functional roles. For instance, network motifs have been shown to form the basis of many complex networks [Milo et al., 2002], especially in brain networks where motif analysis revealed that complex behaviour emerges from local circuit building blocks [Sporns and Kötter, 2004]. Topological network metrics have been used in combination with metadata in some contexts, such as assortative mixing [Bazinet et al., 2023a], but previous studies on network motifs in single neuron resolution data (in *C. elegans* [Varshney et al., 2011, Bentley et al., 2016, Milo et al., 2002] and *Drosophila* [Lin et al., 2024, Hulse et al., 2021]) do not fully exploit the rich metadata available. Comprehensive brain networks are key to understand principles of development [Schlegel et al., 2017], whilst interpreting local connectivity can illuminate the architectural blueprint of the brain [Vogelstein et al., 2019]. In this work, we introduce a method for exploring local connectivity with annotations.

As the number of available connectomes continues to increase, we require more network methods that incorporate the rich complexity of annotations to link the topology of the network to its biological function. Here, we focus on node metadata to introduce a measure of the diversity of local connections and apply this to the *C. elegans* connectome and the *Drosophila* ventral nerve cord (VNC) connectome. The *C. elegans* data set provides the connectome of a complete nervous system, whereas the *Drosophila* nerve cord offers an interpretable model in which information flows between well-defined inputs and outputs with less complexity than the brain [Cheong et al., 2024]. We exploit several different annotation schemes, from fine-grained labels such as cell types to broader categories such as lineages. While there is prior work characterising diversity with respect to community annotations [Bertolero et al., 2017, Guimerà and Amaral, 2005, Guimerà and Nunes Amaral, 2005, Pedersen et al., 2020, Delaplace, 2025, Eagle et al., 2010] and information entropy in network science is not novel [Omar and Plapper, 2020, Bashiri et al., 2020], to our knowledge no prior work aims to characterise the diversity of specialised connectivity in the local neighbourhood of a given node.

In our measure, the *specialisation-diversity*, is derived as follows: first, we construct each neuron’s connectivity vector *p*, which expresses the proportion of incoming or outgoing connections of that neuron to other neurons. The metadata we use are cell classifications or lineage annotations (see **Methods** for details). Then, for a set of connectivity vectors {*p*}, the specialisation-diversity is defined as: 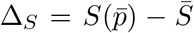 where 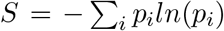, information entropy and 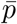 and 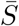 denote the means of {*p*}and *S*. A large Δ_*S*_ value is only possible when the vectors *p* are individually specialised but in diverse categories (see figure 1A and **Methods**). Δ_*S*_ quantifies the level of diverse specialisation in connectivity with respect to the annotations present, and this is a property of the set of connectivity vectors. We consider two sets of vectors for a given neuron that characterise the diversity of its integrative and distributive connectivity. The first is calculated from the diversity of specialisation in the downstream outputs, and the other is calculated from the diversity of specialisation in the upstream inputs. More precisely, for a target neuron, we gather *out*going connectivity vectors of *downstream* partner neurons, and calculate a specialisation-diversity. The resulting Δ_*S*_ associated to this outwards branching motif is what we call the *distributive specialisation-diversity* and notate it as 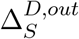. Similarly, a given node’s *upstream* partners’ *in*coming vectors can be used to calculate a specialisation-diversity. We call this *integrative specialisation-diversity*, 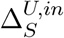. These measures quantify the level of diversely specialised connectivity channels in a node’s local vicinity (figure 1B), with respect to an annotation scheme we use to label the nodes. The annotations we used for the connectivity vectors were cell categories in *C. elegans* and hemilineage in *Drosophila*. These annotations are anatomical and developmental descriptors that are correlated to function and behaviour, and with our measure we are quantifying the most diversely specialised neurons by these associations.

**Figure 1:**
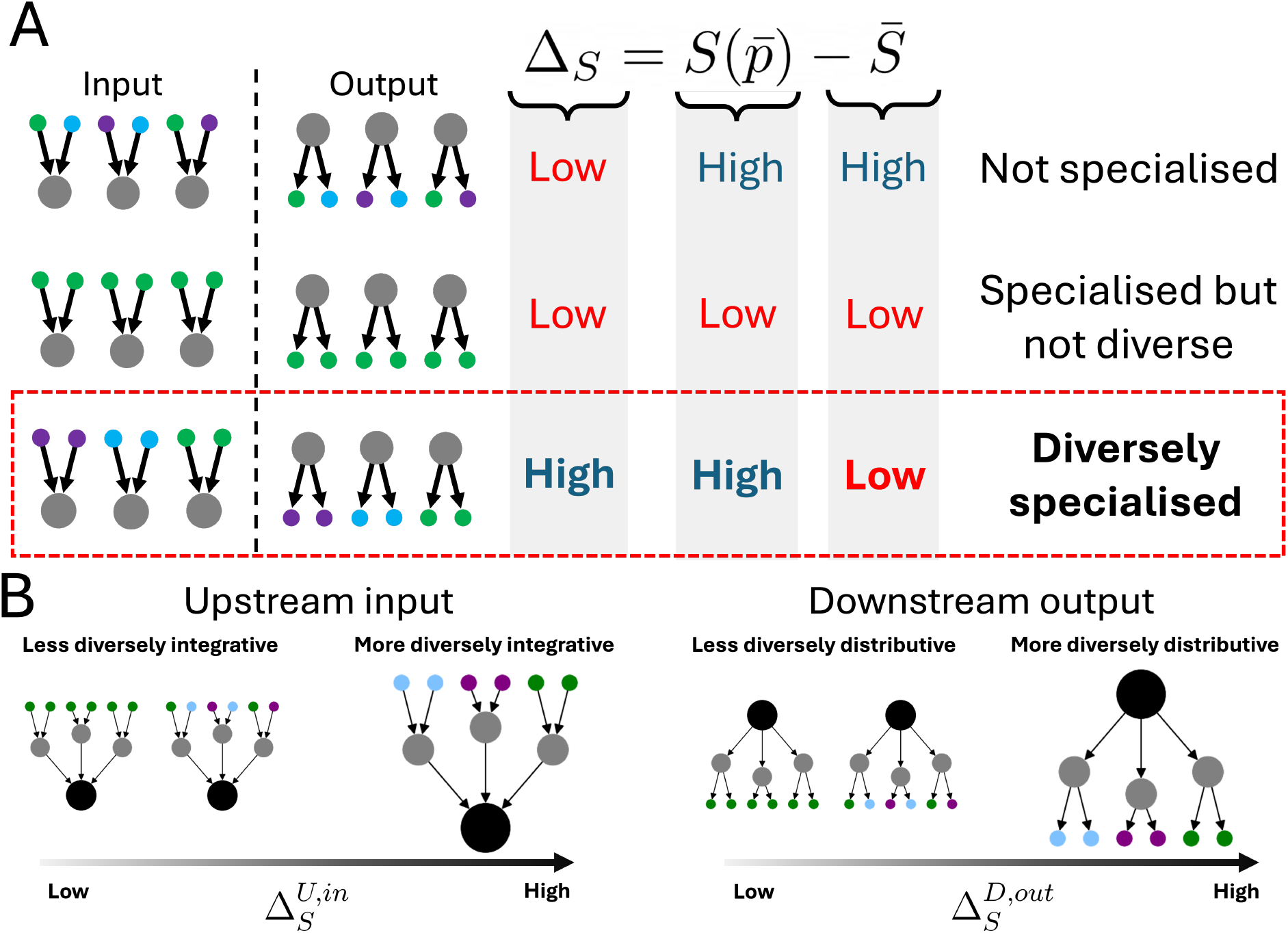
A measure for quantifying diversely specialised connectivity vectors. **A**, Here we illustrate three scenarios concerning input or output connectivity vectors of the three grey nodes (left), where the coloured partners represent different annotation types. The value of entropy is low when the probability distribution is most specialised in one category and highest when it is evenly distributed. In the first row, the individual vectors are not specialised, leading to a high average entropy, 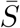and a high entropy of average probability vector 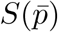. In the middle row, the three grey nodes are specialised in the same annotation type. Therefore, the average entropy is low and the entropy of the average vector is also low (in this case 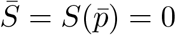). For the above reasons, the two cases yield a low value for Δ_*S*_. In the last row, the grey nodes are also individually specialised in their vectors. In contrast to the previous case, the grey nodes are specialised in different types. Therefore, while the average input entropy is low again, the average input 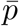 is not specialised and so 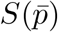 is high. Thus, the difference, Δ_*S*_ is only high when the grey nodes are specialised diversely. We invite readers to the **Supplementary Information**, for a mathematical comment on the bounds of possible Δ_*S*_ values. **B**, We extend the previous argument to two motifs to calculate the specialisation-diversity, and illustrate the cases where the Δ_*S*_ values are low and high. Firstly, with the black node’s upstream partners’ input vectors we measure the integrative specialisation-diversity 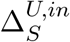 (left). Secondly, the specialisation-diversity of the downstream partners’ outputs measures the distributive specialisation-diversity 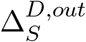 (right).

In this paper, we interpret the diversity of local connectivity as a characteristic of the role of a node in the connectome by using the specialisation-diversity measure. The top 10 neurons with the highest summed integrative and distributive specialisation-diversities are called “entropic hubs”. In *C. elegans* we find these to be linked to multimodal functions having roles in sensorimotor circuits, and in neuropeptide signalling networks [Ripoll-Sánchez et al., 2023]. In the *Drosophila* nerve cord, our entropic hubs are concentrated in the abdominal region where higher-level innate behavioural programmes are controlled [Cheong et al., 2024, Pavlou et al., 2016].

## 2 Results

### 2.1 Cell classes occupy interpretable and distinct regions in specialisation-diversity distributions

We constructed vectors that aggregated the connectivity to the annotations for each neuron. In *Drosophila*, we used hemilineage annotations. These developmental metadata are known to be correlated to certain locomotive functions like walking and flight [Harris et al., 2015]. For *C. elegans*, there have been extensive efforts in cataloguing the neuronal types we may use as metadata. The finest level of classifications were the from gene expression profiles splitting 302 neurons into 118 types [Taylor et al., 2021]. Coarser cell classifications came from identifying broad neuronal function: sensory neuron, motor neuron, pharynx and interneuron classifications. There are ‘mid-scale’ classifications that are based on anatomical position, specific sensory modality or motor programme as well as information flow [Cook et al., 2019]. We chose the mid-scale cell category to construct the vectors that were associated with specialised functions, for instance SN3 labelled neurons are mechanosensory neurons. Additionally, the mid-scale annotations offered a balance such that the connectivity vector would not be too distinct for similar neurons, and conversely, coarse-grained annotations are too general (see **Methods** for a breakdown of neurons into their cell classifications). We reserved the finest and broadest cell type classifications in our validation with null models.

We first verified whether specialisation-diversity was interpretable when considering broad cell classifications in the two connectomes by inspecting the distributions for integrative and distributive diversities (figure 2A for *Drosophila* VNC and 2D for *C. elegans*). In both distributions, neurons are broadly speaking similarly distributive and integrative, but nevertheless there are clear deviations that differ according to the cell classes. In figure 2B and 2E we show the distributions of the quantity 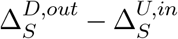 which quantifies how much more distributive or integrative a given neuron is. We report varying biases between the cell types - sensory (orange) being most distributive, and the intrinsic/interneurons (purple) and motor neurons (green) being more integrative. In both *Drosophila* and *C. elegans*, sensory neurons with mostly downstream connections will propagate into the connectome for the processing of external sensory signals, and so are biased towards being distributive rather than integrative. Motor neurons, by contrast, will mostly form upstream connections that control their output, and therefore are more integrative than distributive. The ascending (red) and descending neurons (blue) in *Drosophila* lie in the distributions as we expect: descending neurons being more distributive and ascending more integrative. For intrinsic neurons in *Drosophila* and interneurons in *C. elegans* that mediate the connectivity between the inputs and outputs of the system, we see a slight bias towards integrative diversities. The pharynx (teal) in *C. elegans* is very sparsely connected to the rest of the network and therefore exhibits low distributive and integrative diversities.

**Figure 2:**
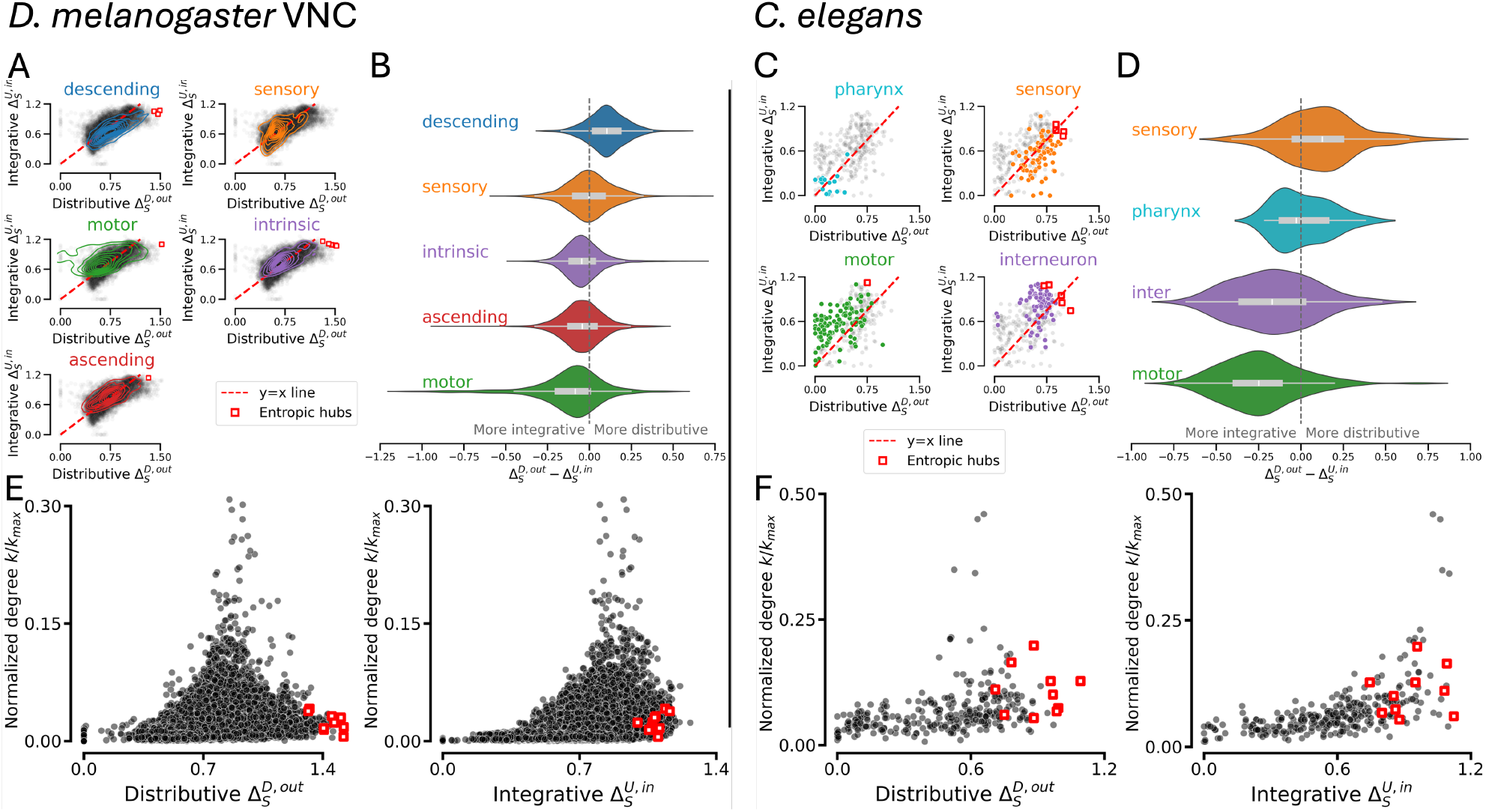
Specialisation-diversity distributions are localised for different cell classes, and are distinct from synaptic degree. The distributions of integrative vs distributive specialisation-diversity for *Drosophila* VNC neurons (**A**) and *C. elegans* (**C**) show localisation of the cell class populations in the plot, with transparent black markers showing the full distribution. In dotted red is the y=x line where the neuron is equally distributive and integrative. Red squares are entropic hubs: neurons that rank in the top 10 highest values of 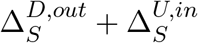. The distribution of distributive minus integrative specialisation-diversity for all neurons in the *Drosophila* VNC (**B**) and *C. elegans* (**E**), reveals that all cell classes have members that are strongly distributive and or integrative, but the majority are well balanced. The averages of the distributions are sorted from most distributive (compared to integrative) at the top to most integrative (compared to distributive) at the bottom. The cell classes are appropriately biased corresponding to their functional role. If the role is primarily projective into the system, then it is more distributive than integrative (descending, sensory). In *Drosophila* the sensory neurons are more balanced as the distribution mean lies around zero, suggesting that the average neuron will be equally integrative and distributive. If the role is primarily integrating diverse channels of information, then outputs in specific and specialised ways, the class is more integrative than distributive (motor, ascending). Intrinsic neurons in *Drosophila* and interneurons in *C. elegans* have a skewed mean towards integrativeness suggesting the function requires more integration of diverse connectivity than distribution. Subfigures **E, F** show that our measure is distinct of degree, the number of total partners of a node. In both animals, the entropic hubness cannot be fully explained by sheer number of synaptic partners, as neurons with the highest specialisation-diversity values occupy low (normalised) degree values.

### 2.2 Entropic hubs are not synaptic hubs

As the distributions of the broad classifications of neurons yield results we can intuitively interpret, we now consider finer classifications and individual neurons. Firstly, we consider the highest-ranking neurons in 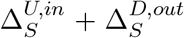 value, which we call *entropic hubs*, (red squares in figure 2). In *Drosophila* the cell classes of the ten highest-ranking entropic hubs are three descending neurons, one ascending neuron, five intrinsic neurons and one motor neuron and all these neurons are more distributive than integrative. These neurons have not been previously described in detail individually, but we note that all of them have soma positions in the abdominal **neuropils**, which are shown to contribute to complex mating behaviour [Pavlou et al., 2016, Marin et al., 2024]. The entropic hubs here are likely all primary (embryonic) neurons (7 labelled, and 3 to be further studied). Primary neurons themselves do not have an associated lineage, but see **Supplementary Information** for analysis of the neurons that have hemilineage information and associated behavioural phenotypes. In *C. elegans*, the entropic hubs play an active role in the neuropeptidergic network (see below). For both connectomes we find that entropic hubs are not synaptic hubs (figure 2C and F), meaning that a high number of synaptic partners does not result in a highly diverse set of upstream or downstream neighbour specialisations. This highlights that our method focuses on the specificity of connectivity rather than the volume of it. The full distributions of all neurons and their metadata are made available (see **Methods**).

### 2.3 Neurons within fine-grained annotation types are similar in terms of their specialisation-diversity values

The finest level of annotation in the connectome data sets are cell types. In *C. elegans* this comprises 118 cell types that are characterised and named according to their morphology and position. In the *Drosophila* VNC, we have cell types that group neurons by identical cell morphology or connectivity. Nervous systems exhibit a well described bilateral left-right symmetry observed in the connectome [Venkatasubramanian and Mann, 2019, Winding et al., 2023, Cheong et al., 2024, Marin et al., 2024]. In *Drosophila* VNC, due to the six-fold symmetry of the leg neuropils, we have six identical analogous neurons forming one *serial homologue* group [Marin et al., 2024, Haszprunar, 1992]. These neurons are annotated by the same label, but are not necessarily strongly connected, so do not form a modular structure within the connectome. Rather, these neurons are similar in morphology so have similar connectivity within each modular structure.

We asked if these finer groups of similar neurons were similar in terms of their specialisation-diversity which was calculated from connectivity aggregation of the mid-scale cell categories for *C. elegans* and hemilineage for *Drosophila*. To answer this, we calculated pairwise distances within the fine groupings using the Euclidean distance on the embedding space of the distributive and integrative diversities [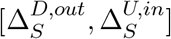]. Explicitly, the Euclidean distance between neurons *i* and *j* is 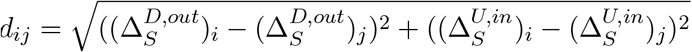. We then tested whether this distribution deviated from appropriate null models (figure 3).

**Figure 3:**
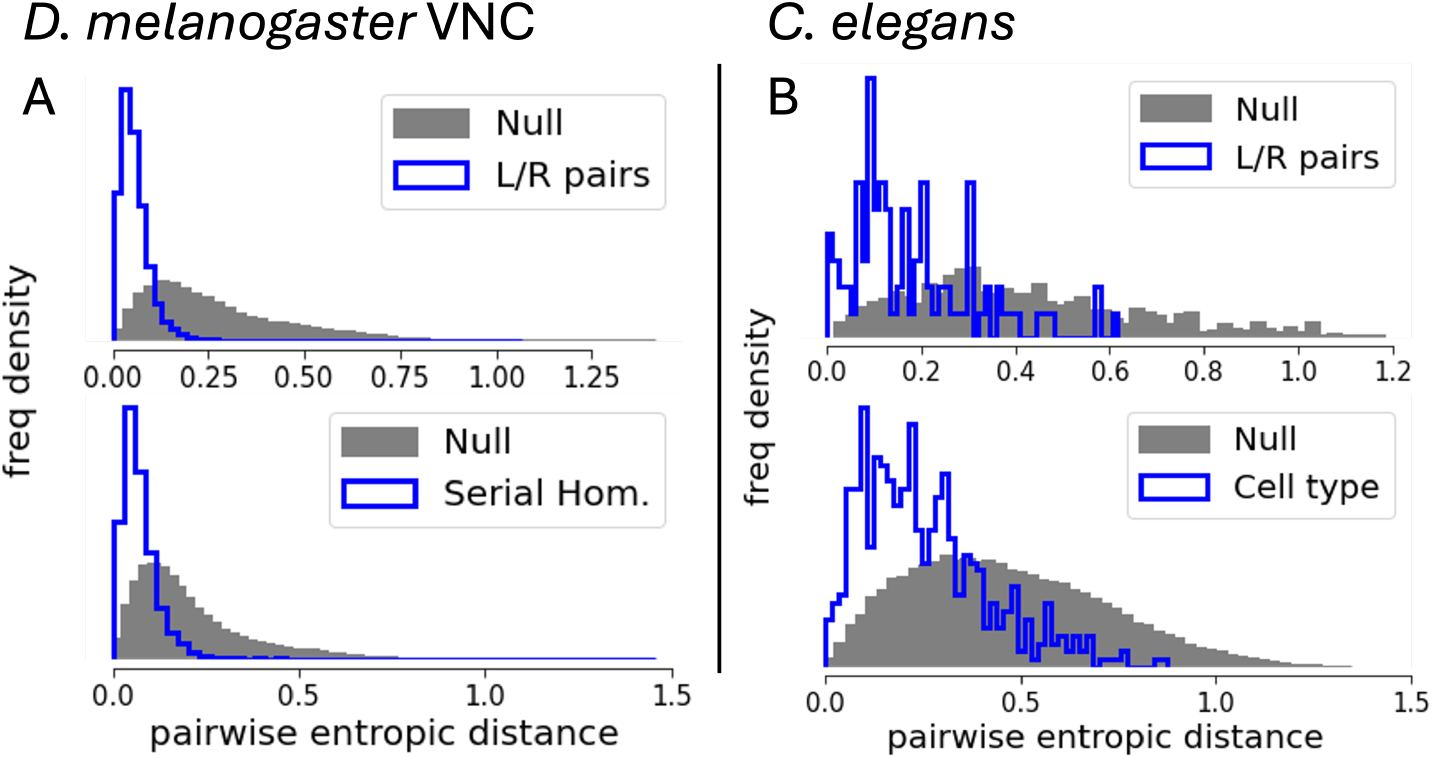
Cell types, left-right pairs and serial homologues are statistically significant in their specialisation-diversity values. The figure shows statistical significance of the specialisation-diversity values for different types of fine grained neuron categorisation. In all cases, we calculate pairwise Euclidean distances of the specialisation-diversity vector 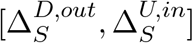, within a specific category and compare this to an appropriate randomised null model. We call this distance the pairwise entropic vector distance. In the VNC (**A**), left-right pairs (top) and serial homologues (bottom) are demonstrated to be statistically significant in their specialisation-diversity values. For left-right pairs, we construct a null model by randomly generating pairs of neurons that conserve the side of the soma, i.e. each randomised pair would be from opposite halves of the VNC. The serial homologue null model picks random groups of six neurons but conserves the leg neuropil membership, i.e. six random neurons from each leg neuropil. In all null models we preserve the connectivity vector and thus the entropy vector. We find qualitatively identical results for *C. elegans* (**B**) using analogous distance calculations using appropriate null models for statistical significance of left-right pairs (top) and fine grained cell types (118 types of which left-right pairs are a subset). The left-right null model again conserves the side membership, and the null model for cell type uses an all-by-all distance calculation (see main text). All null model results are aggregated over 10 realisations of random sampling. Two-sample Kolmogorov-Smirnoff tests yielded significant results for all distribution comparisons with their nulls (*p* << 10^−3^).

We find a statistically significant similarity of contralateral (left-right) pairs using a side-preserving null model of random samples of two neurons, one from each lateral side (top row, figure 3). We also find statistically significant similarity within serial homologue groups in *Drosophila* (figure 3A, bottom), where the null models are pairwise distributions drawn from sampling one neuron from each leg neuropil. Lastly, we tested for pairwise entropic distance within within the 118 cell types in *C. elegans* [Hobert et al., 2016] (figure 3B bottom), with a null model that considers the all-neuron-by-all-neuron pairwise entropic distance distribution. Our results show that similar neurons have similar values of integrative and distributive specialisation-diversities to a statistically significant degree according to a two-sample Kolmogorov-Smirnov test. By testing our null models with the finest-grained cell classifications we have validated our method with the strictest possible criteria.

### 2.4 C. elegans entropic hubs have active roles in complex behaviour

Since all neurons in *C. elegans* are named and have been subject to (in some cases extensive) experimentation, we describe the entropic hubs and their active roles in complex behaviour. The top ranking neuron, `PVR` is a tail interneuron not well described in function. Recently, `PVR` has been identified as a neuropeptidergic network hub, important in the ‘wireless’ neuropeptide-GPCR signalling complementary to the synaptic and gap junction ‘wired’ network [Ripoll-Sánchez et al., 2023]. PVR was also predicted to be a motor control neuron [Yan et al., 2017]. Neuron class `RMG` (`RMGL` and `RMGR`) are interneurons described in the literature as an essential neuropeptide expressor responsible for all types of social behaviour, whose complex behaviour depends on many modes of sensory cues and many diverse repertoire of outputs, with `RMG` being a gap junction hub for connecting 7 classes of neurons [Macosko et al., 2009]. Neuron `DVA` is a stretch sensory neuron for proprioception, and positively and negatively affects locomotion to be able to fine-tune motor neuron stimulation, from a single source. Its central role in regulating the sensory-motor integration explains the diverse downstream specialism [Li et al., 2006]. `DVA` is also an important neuropeptidergic hub [Ripoll-Sánchez et al., 2023] and a hub node linking multiple layers of signalling (synaptic, neuropeptidergic and monoamine) in *C. elegans* Bentley et al. [2016]. The `CEP` neurons are dopaminergic touch sensory neurons in the nose [Kindt et al., 2007, Kang et al., 2010] responsible for food slowing [Sawin et al., 2000] and, along with `ADE`, other behavioural responses to food [Ezcurra et al., 2011, 2016, Chou et al., 2022]. Neuron `URXR` is a body cavity oxygen sensory neuron, central in homeostatic integration of oxygen level fluctuation and metabolic processes. Its sensory signals stimulate body fat loss only when the body has sufficient reserves [Witham et al., 2016]. Neuron `RIGL` is a ring interneuron part of 4 pairs of neurons that mediates CO_2_ response, operating downstream of `BAG` oxygen sensory neurons. Hunger causes the animal to be attracted to CO_2_, and feeding reverses this, and neuron pairs `RIG` and `AIY` mediate this [Rengarajan et al., 2019]. Neuron `SMBDL` is a dorsal neck motor neuron that is a part of the SMB cell type that is crucial in sensorimotor transformation for governing **klinotaxis**, or stimulus provoked lateral locomotion, in particular, changes in salt concentration [Matsumoto et al., 2024].

## 3 Discussion

Our objective in this study was to incorporate biological annotations to enhance our understanding of functional roles in structural brain networks. To achieve this, we have introduced a novel measure based on information entropy that describes the level of diverse specialisation in local directed node neighbourhoods that are integrative and distributive. The specialisation-diversity considers aggregated, directed connectivity from next-nearest network neighbours, thus it is a localised measure for a given node that does not rely on global knowledge.

The application of our approach to the *C. elegans* and *Drosophila* VNC connectomes, using cell classification annotations and hemilineage annotations respectively, yielded distributions of specialisation-diversity that reflect the biological functionalities of these neuron classes. If a given target neuron is on any directed path of length two as the source or sink of that path, then these quantities can be calculated. Therefore, almost all neurons can be quantified by their proximity to specialised connectivity patterns. The capability of neurons to integrate and broadcast the many modes of diversely specialised information aligns with the roles of cell classes. However, all cell classes have outliers that are on either extremes, suggesting heterogeneity and hierarchy in diverse specialisation within class modalities in nervous systems. Interestingly, intrinsic neurons and interneurons that form the bulk of the connectivity bridging inputs and outputs of the network are more integrative than distributive on average. This asymmetry suggests that their connectivity role is to integrate diversely specialised channels of annotation types. Specific entropic hubs with highly integrative and distributive connectivity channels are neurons that are responsible complex sensorimotor behaviour in *C. elegans*, thus requiring close proximity to specialised channels in their inputs and outputs. We propose that entropic hubs in the *Drosophila* VNC are responsible for sensorimotor behaviour, and previous experimental work on the abdominal neuropil suggests an important role in innate behaviours like mating [Pavlou et al., 2016], and so we expect this result to be qualitatively conserved across different individuals.

While we can identify top entropic hubs that are most integrative and distributive, we are limited in terms of the conclusions we can draw from this in *Drosophila*. We have chosen the VNC as a model data set for our analysis, because of its clarity of input and output modalities [Cheong et al., 2024]. With a relative simplicity of sensory and executive brain inputs to motor and feedback into brain outputs, the structure presents an ideal test case that might be difficult to disentangle for more complex data sets, like the brain. We also justify that for the VNC, the hemilineages are strongly linked to motor functions and processing, and analogous conclusions would not be made if similar annotations were not available in other data. The VNC facilitates both reflexive and coordinated movement in the fly independent of brain signalling, the network topology is rich with functional circuitry that allows for complex behaviour [Takemura et al., 2024]. Secondary (post-embryonic) neurons of specific hemilineages have been shown to relate to specific motor functions [Harris et al., 2015], but in our analysis we found the specialisation-diversities to vary between hemilineages annotated with the same function, suggesting further sub-roles, requiring different levels of diversely specialised connectivity (see **Supplementary Information**). Our Δ_*S*_ indirectly measures the neurons that are proximal to many diversely specialised hemilineage connectivity channels responsible for coordinated movement like walking and flight. The entropic hubs in *Drosophila* are mostly primary (embryonic) neurons (7 characterised as such, and 3 not yet annotated). Primary neurons are often globally and centrally connected, and more likely be a member of rich club organisation [Marin et al., 2024], possibly facilitating the high integrative and distributive specialisation-diversities. Early birthtime neurons go through a developmental programme that persists and are “remodelled” during metamorphosis into the adult [Shepherd et al., 2016, Marin et al., 2024] and we speculate that its proximity to highly diversely specialised connectivity are artefacts of dynamic utility and versatility pivotal during development. The entropic hubs are associated with complex, often sensorimotor behaviour in *C. elegans*, with some neurons that were only recently described to be significant like PVR, a neuropeptide hub [Ripoll-Sánchez et al., 2023]. We also report that entropic hubs are not conventional hubs in terms of their degree in the synaptic network. In *C. elegans*, while some entropic hubs have high degrees compared to the rest of the network, there are several neurons that exhibit highly distributive and integrative diversities without being synaptic hubs. In *Drosophila*, the absence of correlation between specialisation-diversity and degree is clearer. We observe that while highly connected nodes could also be in close proximity to highly specialised and diverse connectivity channels, their high degree makes them more likely to be connected to the same specialisations, and non-specialised partners, which penalises their specialisation-diversity value. This is consistent with previous studies that showed that node degree was independent of a node’s diversity of connection to disparate modules [Pedersen et al., 2020].

We have verified that the cell types, contralateral pairs and serial homologues are more similar to each other in specialisation-diversity values than randomised null models. Thus, similar neurons are also similarly diversely connected. This is expected as cell types are often defined to be an irreducible subunit of neuronal grouping [Dance, 2024, Schlegel et al., 2024, Hobert et al., 2016]. Therefore, they must be functionally identical and so identically connected. The statistical significance of the pairwise entropic distances against various nulls of cell types and serial homologues contributes to providing comparative methodologies for cross-identifying and contrasting different connectome data sets [Schlegel et al., 2024, Stürner et al., 2024]. However, the specialisation-diversity values of functionally different neurons overlap considerably. While we are able to recover the broad cell classifications to be localised at low or high integrative or distributive specialisation-diversity values, it is insufficient to infer the specific finer grained categorisations from specialisation-diversity alone. Since we have reduced the network of annotated nodes into the two numbers quantifying the proximity to diversely specialised channels of connectivity, equivalently diverse connectivity may arise from different connectivity specialisation. We leave this inverse inference problem for future work, besides approaches like graph neural networks [Corso et al., 2024] have demonstrated convincing evidence that node annotations supplement our inference of function in the underlying network. We also speculate more open questions about the nature of biological annotations complementing the network connectivity. Many different annotation schemes are available from coarse to fine-grained; node or edge metadata and their informative or predictive relationships with network structure are not fully understood [Bazinet et al., 2023a,b]. While our methodology can be extended to handle these, we do not yet know of which annotation schemes are most biologically and inferentially complementary for detecting specific functional attributes of the underlying system.

The connectomes we have used here also have limitations, for instance in terms of the incompleteness of annotations in *Drosophila*. Hemilineage annotations are inferred from morphology and connectivity and are incomplete for primary (embryonic) neurons which are functionally and morphologically distinct from secondary (post-embryonic) neurons. We ignore connectivity with unlabelled nodes in our methodology, but expect our conclusions to remain qualitatively robust since primary neurons arguably should be treated as a separate hemilineage subtype in our methodology because their functions and morphology are distinct to secondary neurons of the same lineage. Hemilineages are developmentally driven functional units of the nerve cord, thus we expect the average connectivity statistics to be conserved across individuals, as demonstrated in recent comparative studies [Lin et al., 2024, Stürner et al., 2024, Schlegel et al., 2024]. Moreover, the *C. elegans* connectome has been shown high connectivity conservation across individuals and developmental stages [Cook et al., 2019, Witvliet et al., 2021], thus we expect our results to conform on average with new data. Another limitation of the work is that our results are speculative and predictive rather than conclusive of function or activity which has been described in anatomical perspectives of neuroscience previously [Prinz et al., 2004]. Since our specialisation-diversity pools connectivity from all neurons that are reachable by or with a directed path of length two, we are compounding biological variability, or errors attributed to connectome imaging and reconstruction present in the data. Despite this, our analysis shows sufficient evidence that aggregating by connectivity produces statistically significant results to justify this study, and so expect our conclusions be qualitatively similar with future data that arises from advances in the acquisition of connectomic data. We expect future connectome data sets to be accompanied by a variety of annotations, from morphology to genetic annotations. Annotations are likely to be a key aim of synapse resolution connectomics pipelines to catalogue the neural inventory that informs other neuroscientific fields. This paper serves as a proof of principle of our method that is shown to provide interpretable insights in connectomes across two different magnitudes. This indicates that our specialisation-diversity can be straightforwardly applied to larger and more complex data sets, like the full *Drosophila* brain [Dorkenwald et al., 2024].

Our measure aggregates the connection profiles of neurons that are reachable by a path of length two from a given query neuron, and then compares their specialisation. There are opportunities to extend this measure for interesting metrics quantifying the diversity of connectivity. The base methodology can take any set of normalised probability distributions to quantify the diversity of the specialisation. An opportunity for future work exists to extend the method to consider specific circuits in the network and calculate the specialisation-diversity of neurons along a path. These path-based measures could quantify the distinct annotation profiles that could exist in a query signalling pathway, where the extension of the metric may allow comparisons of the extent of diverse integration or distribution of these query pathways. Additionally, inspecting the specific annotations that contribute to the pathway scoring highly in a specialisation-diversity measure may elucidate the specific modalities contributing to the information processing along the path. For instance a touch-sensory to a body motor neuron pathway may exhibit a repelling or attracting behaviour depending on other sensory cues.

Data on the large-scale organisation of nervous systems has only recently become available to the neuroscientific community. Our primary aim was to provide an augmented perspective of network analysis that incorporates node annotations. To do this, we introduced an entropy-based measure, demonstrated its potential as an interpretable descriptor of neuronal function, and proposed roles for neurons not yet individually characterised experimentally. More generally specialisation-diversity is a local centrality measure in annotated networks, that can propose key nodes that are important for aggregating and disseminating diversely specialised channels of connectivity. Similarly, this structural identification of the possible integration and distribution of annotation connectivity can be expanded to other connectome types. With a careful choice of annotations the method could reveal diversely integrative and distributive regions. Nodes with diverse connections but not necessarily large numbers of connections are especially interesting in neuroscience [Bertolero et al., 2017, Pedersen et al., 2020], and our methodology requires high coverage of meaningful annotations to uncover these structures. Other annotations that are useful might be transcriptomic profiles that catalogue the molecular functionality in each region, ideal for meso-scale connectomes without general single-neuron knowledge to characterise functionality. However, we do not expect to exactly retrieve our results for networks of different resolution, as by construction, the complexity of connectivity which our method relies on is averaged out. While we have demonstrated our method using neuronal connectomes, it can therefore be applied to any complex network with node annotations.

## 4 Methods

### 4.1 Specialisation-diversity

To calculate the specialisation-diversity, we require normalised connectivity vectors describing the proportion of connectivity of a given neuron to each annotation label. We first establish the labels for all nodes we adhere to for connectivity aggregation. For given target node, we collect its partners whose labels are known. The frequency distribution of the node label types that are connected to the target node is then determined. We normalise by dividing by the total in or out strength of each frequency of node label, giving the discrete probability distribution of the in/out-connections to each of the labels. This removes biases of scale in the network, and addresses the in and out connections separately. The resulting vector has entries corresponding to proportion of contributions *in*-from or *out* -to each node label type.

Mathematically, let *a*_*iz*_ be the elements of an adjacency matrix, denoting the weight of a directed edge from neuron *i* to neuron *z*, with value zero if the edge does not exist. Let *b*_*zj*_ be another matrix element where *b*_*zj*_ = 1 if node *z* is labelled with annotation type *x*_*j*_, (e.g. a specific hemilineage) and zero otherwise. Then the vector of annotation type contributions for the in and out connectivity becomes:

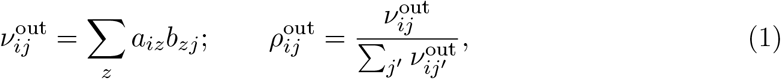

and the in-vector is the same, with a transposed adjacency matrix element (*a*_*zi*_):

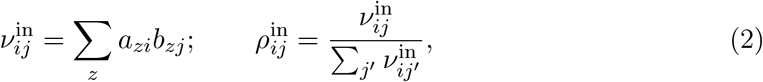

where the probability vectors (*ρ*) are normalised to sum to unity, and can write as 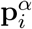, where *α* ∈ [in, out] can denote an in-vector or out-vector in reference to node *i*. For clarity, let’s drop *α* as the derivations are identical and discuss in and out directions later.

Notice that **p**_*i*_ is a probability distribution equivalent to *p*^(*i*)^(*X*) where *X* is a random variable that can take the values of the annotation types, *X* ∈ {*x*_1_, *x*_2_, …, *x*_*M*_ }. We use the shorthand for probability for annotation type *x*_*j*_ as 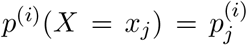. Then, by using information entropy, we can measure specialisation in this particular probability distribution:

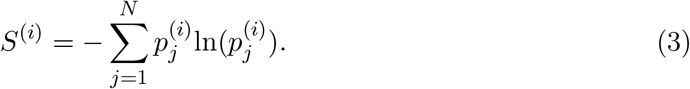

Entropy is maximised when all contributions *p*_*j*_ are equal, meaning the node is connected to all types of labels equally so it is not specialised. Entropy is minimal when the connectivity is concentrated rather than distributed, with the minimum entropy value being 0 where all connections are with one label type. This means that specialised nodes have low entropy.

Now, if we have many probability vectors, we can calculate how specialised they are individually and so compare how diversely specialised the probability vectors are. Let *P* = { *p*^(1)^(*X*), *p*^(2)^(*X*), …, *p*^(*N*)^(*X*)} be a set of normalised connectivity probability distributions, where *N* is the number of distributions. Associated with the probability vectors, are each of their individual entropies: *S* = {*s*^(1)^, *s*^(2)^, …, *s*^(*N*)^ }.

The average entropy is given by:

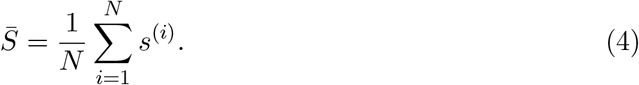

The average connectivity vector component 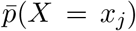, written with a shorthand 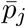 is given by:

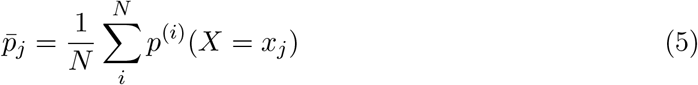

which can be demonstrated that it is also a probability distribution with the property 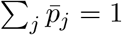 (see **Supplementary Information**). Thus, the entropy of the average vector can be calculated and meaningfully compared:

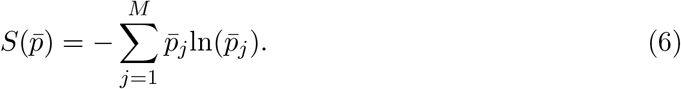

We are particularly interested in neurons are connected to other neurons with their own specific specialisms which are individually diverse. If a given neuron’s connectivity is concentrated or specialised, then the entropy of this neuron will be low. If a group of neurons whose connectivity is specialised in the same node label types, then the average entropy, 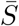, will be low as well as the entropy of the average vector, 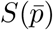. If a group of neurons whose connectivity is specialised in different node label types, the average entropy, 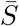, will be low, however, the average vector will not be specialised so the entropy of the average vector, 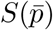, will be high. This is illustrated in figure 1.

If the average entropy is high, the group of nodes are unspecialised so the average vector must also be unspecialised and the entropy of the average vector is also high. With this, we observe that large differences between the average entropy and the entropy of the average vector mean that the group of neurons have specialisms in a variety of different node label types. If the entropy averages’ differences are small, then it could mean that the group of neurons were not specialised to begin with, or that the group of neurons were all specialised in similar node label types (figure 1A, top two rows).

Using these measures, we demonstrate that a high difference in entropy of the average vector and the average entropy of each vector is only observed when the set of vectors are diversely specialised (figure 1A, bottom row). We arrive at the Δ_*S*_, the *average entropies’ differences* - the difference in the average entropy and the entropy of the average vector, which measures the *specialisation-diversity* :

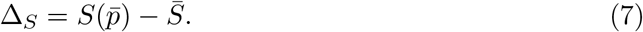

Entropy is a concave function, so negative entropy is a convex function. This means that negative entropy must obey Jensen’s inequality for convex functions: 𝔼 [*ϕ*(*X*)]−*ϕ*(𝔼 (*X*)) ≥ 0, where 𝔼 denotes the expected value and *ϕ* is a convex function on a probability distribution *X*. Therefore, substituting for −*S*(*p*) = *ϕ*(*X*), and ⨪ for the expected value, 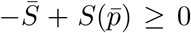, and so Δ_*S*_ ≥ 0 always. This is logically sound, as a set of probability vectors would not become more specialised than the individual constituent vectors through averaging. In other words, the entropy of the average vector would always be at least value of the average entropy of those vectors.

Notice that have two choices of the set of probability distributions *P* we wish to calculate the specialisation-diversity: which set of nodes *i* = 0, …, *N* to consider, and which direction the connectivity is aggregated (*α* ∈ [in, out] we dropped earlier). We are free to calculate a specialisation-diversity for any combinations of upstream or downstream partners; input or output vectors. We focus here only on the upstream inputs (integrative) and downstream outputs (distributive), and label them 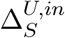and 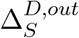 respectively. In other words, for a given target node, we may collect its upstream partners’ input vectors and we can calculate the specialisation-diversity associated to this (figure 1B, right). Secondly, we may use the downstream partners’ output vectors of a given node to calculate the specialisation-diversity.

### 4.2 Drosophila VNC and C. elegans data sets

The *Drosophila* VNC connectome and associated annotations used in this paper is the Male Adult Nerve Cord (MANC) data available from https://neuprint.janelia.org/publically available with a Google account (see Takemura et al. [2024]). The *C. elegans* connectome is a hermaphrodite data set from www.nemanode.org [Witvliet et al., 2021] by Varshney et al. [2011] (based on White et al. [1986]) and the associated metadata was compiled from Cook et al. [2019] available from https://www.wormatlas.org/neuronalwiring.html, and Ripoll-Sánchez et al. [2023].

Annotations that were used in this paper for the connectivity vectorisation for specialisation-diversity calculations were available from the data sources listed above. For *Drosophila*, we have hemilineage determined from light level data and morphology and connectivity clustering detailed in Marin et al. [2024]. The secondary neurons of hemilineages have been experimentally demonstrated to form behavioural phenotypes in independent of brain signalling [Harris et al., 2015]. Here, lineages are specified by neuronal precursor cells (ganglion mother cell) which then diverge via Notch signalling into two distinct phenotypes ‘A’ and ‘B’ which form the hemilineages. These hemilineages are the developmental units of organisation in *Drosophila* [Truman et al., 2010]. By calculating the specialisation-diversities with the hemilineage annotation scheme, we are interested in how diversely specialised are the connectivity channels that perform these functions. In the **Supplementary Information**, we investigate whether specialisation-diversity averaged for each hemilineage reveals neurons that are highly important as they are proximal for collecting information from upstream and dispersing information downstream of many different functions.

In *C. elegans* the annotations we use are mid-scale cell categorisation by Cook et al. [2019], based on anatomy and information flow. This further splits interneurons into layers, specifying the sensory modality and motor location. These labels are directly associated with functionality, as opposed to the high-level cell classes (motor, sensory, interneuron and pharynx). The other alternative was to use the 118 cell types determined from transcriptomic profiles [Taylor et al., 2021]. We deemed the high-level cell classes to be too broad, and the transcriptomic classes to be too specific. If the annotations are too fine, then the connectivity profiles may be too dissimilar between similar neurons; conversely, coarse-grained annotations are too general so the neighbourhood vectors are too similar. Therefore, we utilised the mid-scale cell categories for constructing our annotation vectors. We implemented all null distribution tests with the finest cell types for the strictest test of correlation to known neurobiological criteria. These cell types are presented in table 2 (**Supplementary Information**). We tested alternative annotations (fine and coarse) in the vector construction for the specialisation-diversity calculation, but did not offer as interpretable results for the above reasons. Exact determination of how optimal an annotation scheme is correlated to what interpretable property of the system is an open question.

### 4.3 Code availability

All files for python analysis for generating figures is documented and made available from https://github.com/sungsushi/spec_div. We include all data and processed data from this work in https://doi.org/10.5281/zenodo.15052465.

We utilise the scipy.stats.ks_2samp implementation of the two-sample Kolmogorov-Smirnov test.

## Acknowledgments

We thank Gregory S. X. E. Jefferis, Philipp Schlegel, Tomke Stürner and Elizabeth Marin for discussions and providing access to data.

S.S.M. acknowledges the support of the Harding Distinguished Postgraduate Scholars Programme Leverage Scheme and the EPSRC DTP.

## Supplementary Information

### Proof that the category-wise mean of a set of discrete probability distributions is also a probability distribution

Let *P* = {*p*^(1)^(*X*), *p*^(2)^(*X*), …, *p*^(*N*)^(*X*) } be a set of probability distributions, where *N* is the number of distributions and *x* is the discrete random variable *X*∈ { *x*_1_, *x*_2_, …, *x*_*M*_ }.

By definition, the sum of the *i*th probability distribution adds to one:

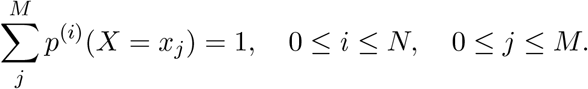

We take the category-wise mean of this set of probability distributions:

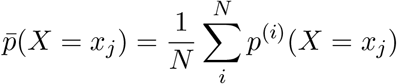

which is the probability of *x* averaged over all the distributions. 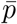 must be positive and bounded by the interval [0, 1] since the constituent probability distributions are also positive and bounded for all *i, j*.

To check that this distribution indeed does sum to unity:

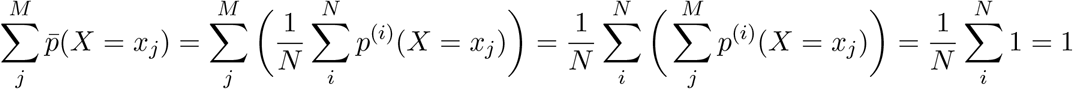

by using the previous definition of the probability vectors. Therefore, the category-wise mean of a set of discrete probability distributions is also a probability distribution.

### Alternate form for Δ_*S*_

Here, we derive an alternate form for the specialisation-diversity which is helpful for implementation purposes and finding bounds on the metric.

From the definitions of 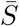, 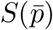 and Δ_*S*_ in equations 4, 6 and 7:

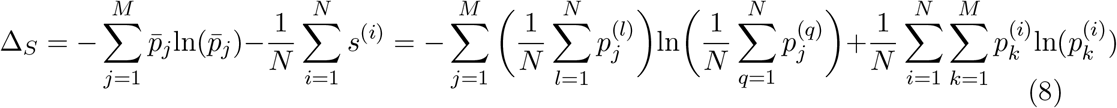

The first term can be simplified as follows:

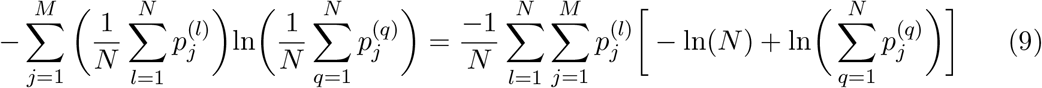

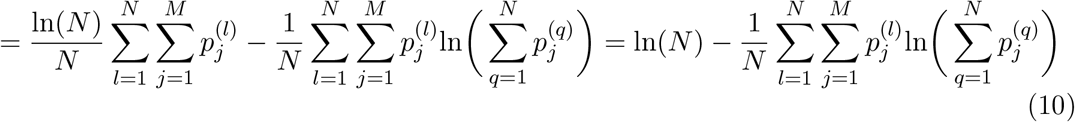

where we invoke that the category-wise average of a probability distribution is itself a probability distribution and must sum to one.

Now, substituting this back to equation 8 :

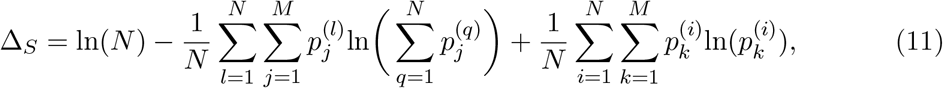

we may rewrite the sums over the labels and the sums over the vectors (i.e. setting *j* = *k*

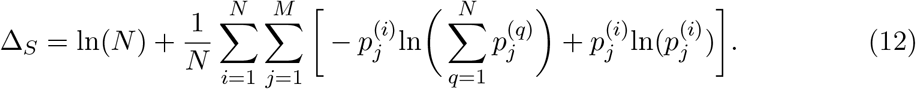

Then, writing it as one logarithm:

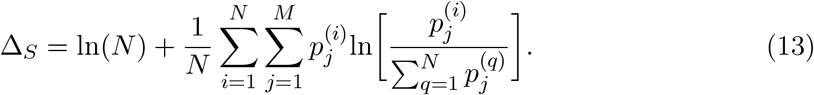

We consider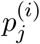 to be the probability of label *j* given that the distribution is the *i*th vector, and can write as a conditional probability: *p*(*j*| *i*). And so in this interpretation, the term

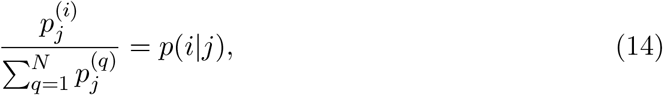

as it is the probability of choosing vector *i* given that we consider label *j*. The notation is consistent with conditional probability obeying Bayes’ rule.

Finally, we arrive at:

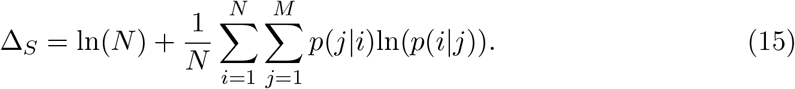

### Bounds on Δ_*S*_

The lower bound for specialisation-diversity is zero due to Jensen’s inequality. For an upper bound, consider equation 15. The second term is always negative since *p*(*j*|*i*)ln(*p*(*i*|*j*)) ≤ 0 as 0 ≤ *p*(*i*| *j*) ≤ 1. (We make use of the limit lim_*x*→0_ *x*ln(*x*) → 0, and by definition in equation 14, *p*(*j*| *i*) = 0 if *p*(*i*| *j*) = 0.) Consider the case where we have fewer vectors than categories, *N < M*. Δ_*S*_ is maximized when *p*(*j*| *i*)ln(*p*(*i*| *j*)) = 0 for all pairs of *i, j*. Physically, this would mean maximally specialised (*p*(*i*| *j*) = *p*(*j*| *i*) = 1) in different categories, and the remaining entries being zero (*p*(*i*| *j*) = *p*(*j*| *i*) = 0). And so the maximum value for Δ_*S*_ is ln(*N*). When there are more vectors than categories (*N > M*), we must have a degeneracy in specialism even in the most diversely specialised scenario, thus imposes a stricter bound Δ_*S*_ *<* ln(*N*). This would contribute to degree hubs (large *N* compared to *M* , the number of annotation types) being penalized in their specialisation-diversity value if their partners are degenerately specialised.

### Hemilineage functionality is reflected in specialisation-diversity distributions

From Harris et al. [2015], we know which *Drosophila* hemilineages (secondary neurons) activate when certain functions are observed. These are: changes in posture, uncoordinated leg movement, walking, wing wave, wing buzz and take-off, and many hemilineages were activiated in more than one functions. We binarized these relationships, and sought to ask are the complexities in functionality, which is summarized in table 1.

**Table 1:**
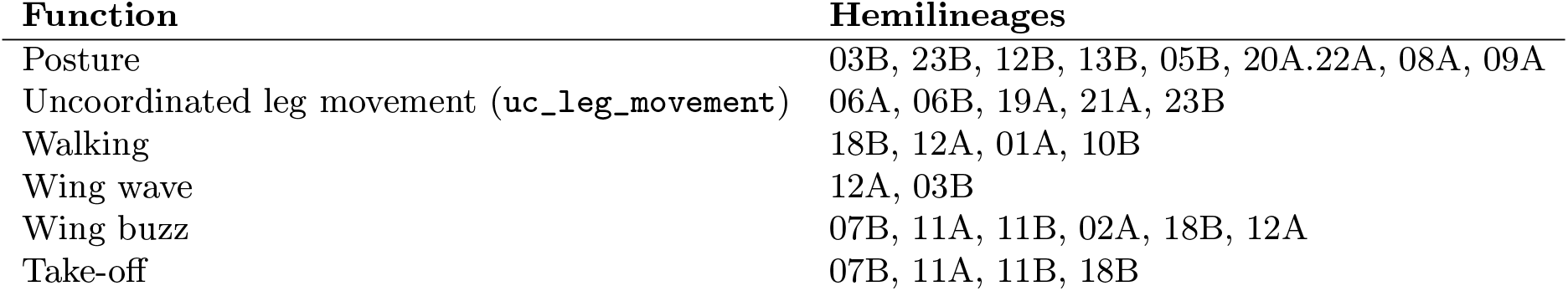
Hemilineages and their link to motor functions in the VNC in the absence of brain signalling from Harris et al. [2015].

When we averaged the integrative and distributive specialisation-diversities we found that even with the ‘same’ functional were very different in specialisation-diversity values (figure 4), suggesting that the functional categorisation may offer itself to further sub-roles that require different levels of diversely specialised connectivity channels.

**Figure 4:**
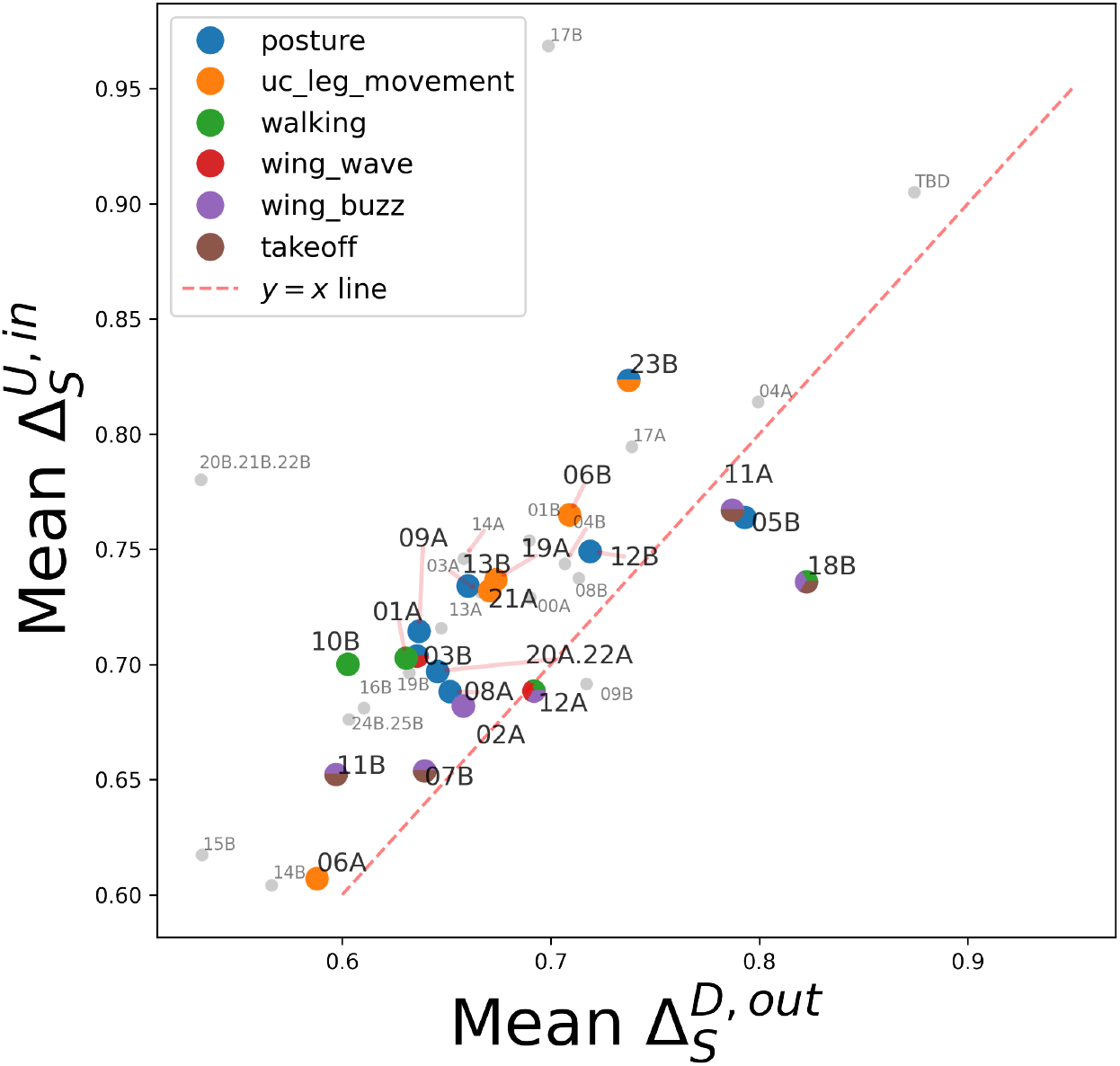
Hemilineages with same function may have more specialised roles. Here, we plot the mean integrative vs distributive specialisation-diversities averaged across all secondary birthtime (post-embryonic) neurons of each hemilineage in the *Drosophila* VNC. We note each function by six colours in the legend, and split the marker colours when a hemilineage is associated to multiple functions. In grey are hemilineages not described in Harris et al. [2015]. The red dotted *y* = *x* line is where points are equally distributive and integrative.

## 5 Table of cell classes in *C. elegans*

We show the coarse, mid-scale and fine annotations associated with the *C. elegans* neurons in table 2, see **Methods** for details.

**Table 2:**
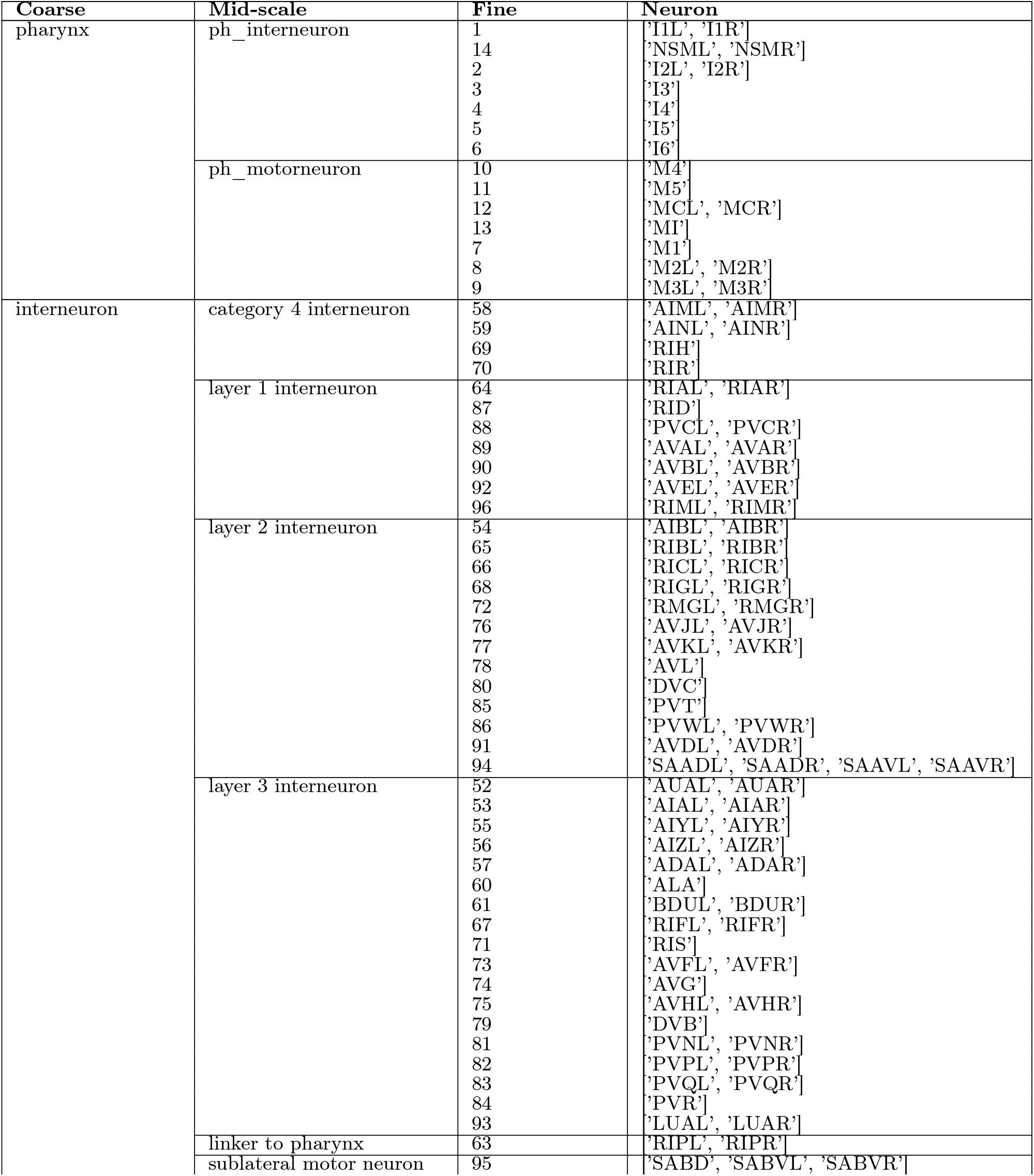
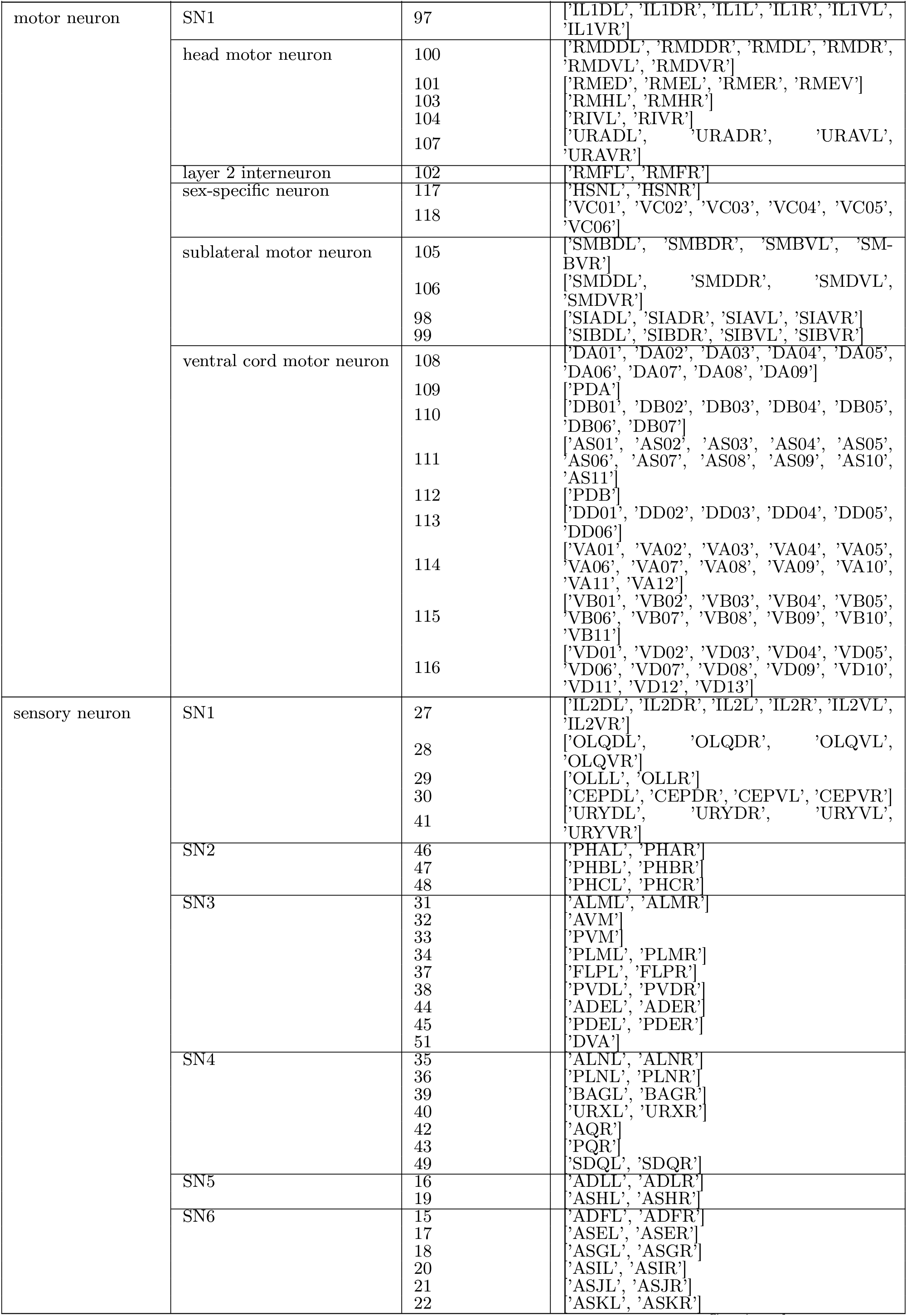
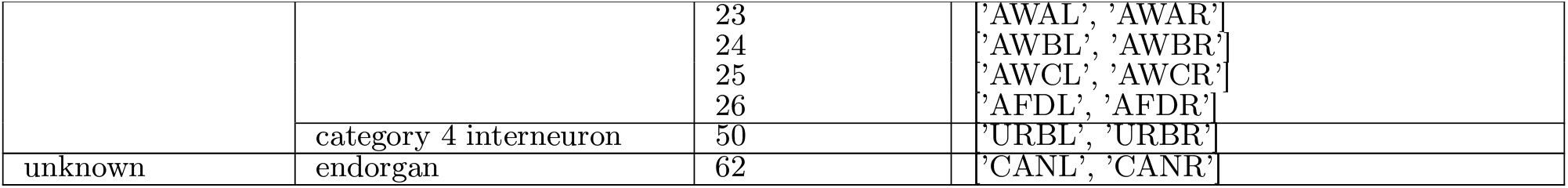
Three neuronal classifications of various coarseness are given. The coarsest cell class is from Ripoll-Sánchez et al. [2023], “Final Classification”, the intermediary mid-scale cell category used to construct the annotation vectors is from Cook et al. [2019] under the name “cell category” and the finest types are the canonical 118 numerical “cell class” labels from Ripoll-Sánchez et al. [2023], compiled from morphological, transcriptomic and connectivity characterisation Taylor et al. [2021], Hobert et al. [2016].

